# How to prevent chick culling in the poultry industry? Discovery of a new biomarker for *in ovo* gender screening

**DOI:** 10.1101/2023.08.20.551526

**Authors:** Nicolas Drouin, Hyung Lim Elfrink, Wouter Bruins, Slavik Koval, Amy C Harms, Wil Stutterheim, Thomas Hankemeier

**Affiliations:** Metabolomics and Analytics Centre, Leiden Academic Centre for Drug Research, Leiden University, Einstein weg 55 2333CC, Leiden, the Netherlands; In Ovo, Haagse Schouwweg 12, 2332 KG, Leiden, the Netherlands

**Author notes:** **Corresponding Author:** Prof. Thomas Hankemeier. equal contribution (Nicolas Drouin, Hyung Lim Elfrink, Wouter Bruins).

**Keywords:** Mass spectrometry, metabolomics, biomarker, gender screening, chicken eggs, animal welfare, poultry industry

## Abstract

Chicken eggs are one of the most consumed foods worldwide. However, the practice of chicken culling in the poultry industry involves unnecessary animal suffering and finding a way to put an end to this has become a societal priority. One approach that has been propagated as acceptable is based on the selection of female eggs early in the incubation process and the devitalization of the male eggs.

It is with this objective in mind that we searched for a biomarker for early gender screening in eggs. Applying an untargeted mass spectrometry approach, we profiled allantoic fluid of different day-old eggs and identified the feature 3-[(2-aminoethyl)sulfanyl]butanoic acid (ASBA) as a strong biomarker for *in-ovo* gender prediction for day-9 old embryos. After validation using LC-APCI-MRM with an internal standard, we found ASBA can predict the female gender with a sensitivity and specificity well above 95% in our experiments.

**Highlights:**

- Discovery of a biomarker of chicken embryo gender in allantoic fluid from eggs
- Day 9 after laying was determined as optimum for sex prediction and animal welfare
- 3-[(2-aminoethyl)sulfanyl]butanoic acid was identified using mass spectrometry
- The biomarker was validated on large cohorts of different chicken species

**Graphical abstract:** 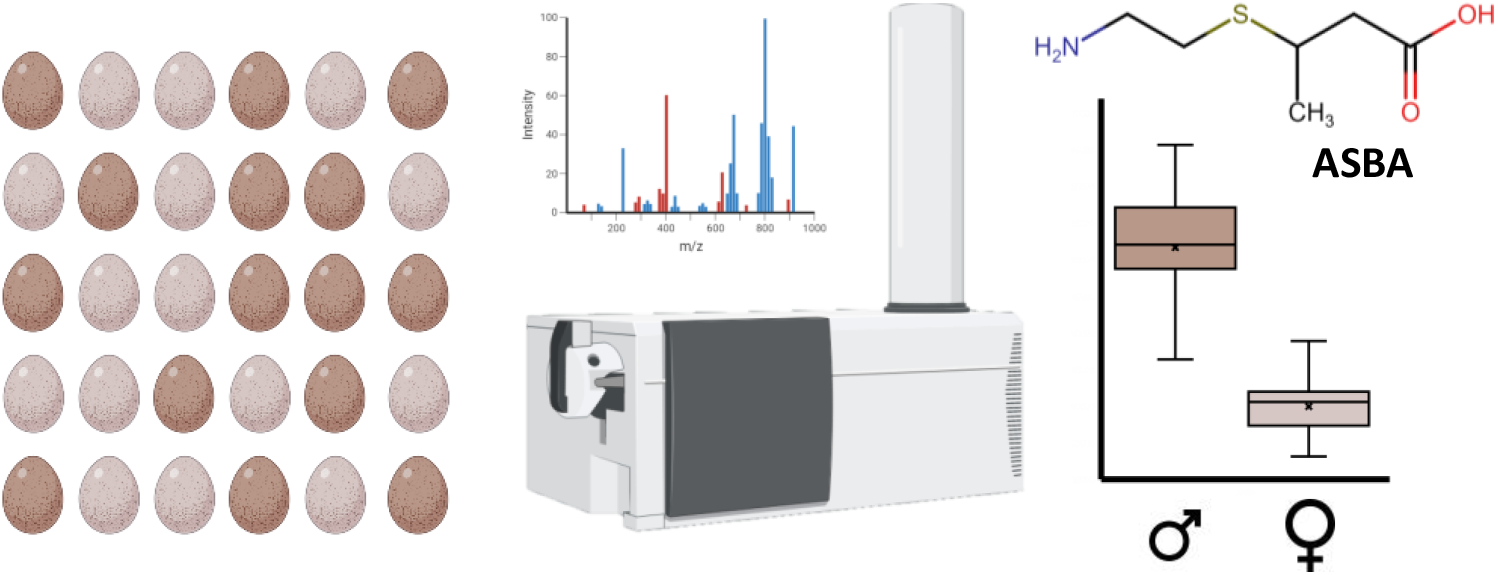

## 1. Introduction

Chicken eggs are one of the most popular cooking ingredients in every country over the world. Every year, more than 1.3 trillion eggs are produced by 4.7-6.5 billion of laying hens, representing a tremendous 80.1 million metric tons of eggs (WATTPoultry, 2019). The productivity of laying hens drops anywhere between 1 to 3 years of age after which they are slaughtered and replaced. This causes a perpetual need for massive laying hen production. Unlike broiler chicken breeds, males from these egg-laying breeds do not produce enough meat to have any significant value in the food chain. Hatching males increases the costs in the laying hen industry. Furthermore, this poses an ethical problem: the need to cull the males. More than 3.2 billion day-old male chickens are culled every year. Currently, due to lack of viable alternatives, chick sexing is manually performed after hatching and, depending on the breed, is based on different colors of the down, or observation of the cloaca. While female chicks are kept for egg production, males are killed by mechanical methods such as maceration, electrocution or asphyxiation (Health, Welfare, Nielsen, Alvarez, Bicout, Calistri, et al., 2019; UNION, 2009).

Interest in animal welfare has been growing over the last decade. Therefore, several countries, such as France, Germany, Switzerland or USA, are currently designing new and more restricted legislation on this practice in the poultry industry (Blide & Montel, 2015; Commion de la Science 2019; l’Alimentation, 2020; Producers, 2020). This adds to the urgency to find economically viable solutions to end male chick culling. In this context, *in ovo* gender determination to select and sort eggs before hatching is one of the most promising strategies. In the early stages of the embryo development, the central nervous system is not fully developed. Consequently, the embryo is not able to feel the sensation of pain (Close, Banister, Baumans, Bernoth, Bromage, Bunyan, et al., 1997). This would therefore make the early removal of male eggs a more acceptable practice.

As any gendered species, gender can be determined with absolute certainty by genotyping. In that respect, He *et al*. developed sensitive and robust PCR and qPCR approaches (He, Martins, Huguenin, Van, Manso, Galindo, et al., 2019). However, PCR based approaches are expensive and too slow to screen each egg in a hatchery. Some alternative methods have already been developed for this purpose. In 2011, Steiner *et al*. developed a FT-IR spectroscopy method to detect the chromosomic difference between males and females from the germinal disc (Steiner, Bartels, Stelling, Krautwald-Junghanns, Fuhrmann, Sablinskas, et al., 2011). However, this method required puncture and collection of cells from the embryo, presenting a high risk for the fetus. More recently, the same group developed Raman and fluorescence spectroscopy methods targeting the blood vessels located under the shell (Galli, Preusse, Uckermann, Bartels, Krautwald-Junghanns, Koch, et al., 2016, 2017). The combination of both spectroscopic measurements attained an overall correct prediction rate of 93% in small-scale experiments (Galli, Preusse, Schnabel, Bartels, Cramer, Krautwald-Junghanns, et al., 2018). Only a handful of processes have reached a pilot test scale. Among these figures the method developed by Weissmann *et al*., which is now allowing for a limited commercial production by Seleggt GmbH. They identified estrone sulfate, a metabolism product of sex steroids, as a biomarker for *in ovo* early gender prediction and they developed an ELISA test with a sensitivity of 86.0% and specificity of 82.9% (Weissmann, Reitemeier, Hahn, Gottschalk, & Einspanier, 2013). However, immunoassays are expensive due to the manufacture of selective antibodies and are relatively low throughput that makes this method largely unsuitable for a large-scale industrial application. Also, the method can only be applied at a relatively late stage of the embryonic development due to the late occurrence of estrogen sulfate in the embryo development.

The present study reports the discovery and characterization of a new biomarker using mass spectrometry-based analytical tools. The identification of the relevant features was first made using untargeted RPLC-MS analysis and subsequent structure elucidation of these features was achieved by the combination of several analytical approaches using MS/MS. The newly discovered biomarker allowed an early female gender determination at day-9 in the experiments described below. We also demonstrate its prediction capability in different chicken breeds.

## 2. Materials and methods

### 2.1. Chemicals

S-Propyl-L-Cysteine and 3-[(2-aminoethyl)sulfanyl]butanoic acid (ASBA) and the deuterated version were synthesized upon request by Enamine (Kiev, Ukraine). S-(2-carboxypropyl)-Cysteamine was synthetized by the group of Prof. Byeong-Seon Jeong (Gyeongsan, South Korea). MS grade water was produced by a milliQ water generator. Formic acid, methanol and acetonitrile were purchased from Acros Organics (Geel, Belgium) and Actu-All Chemicals (Oss, the Netherlands) respectively.

### 2.2. Samples

All eggs used in this study came from brown chickens (VB1636 Brown Nick) and white chickens (Hy-Line CV 24). Depending on the development stage, from 100 μL to 1 ml of allantoic fluid has been manually collected by carefully drilling a hole in the top of the shell. A 0.22 Gauge needle on a syringe was used to extract the allantoic fluid without inflicting harm to the developing embryo. After collection, the allantoic fluid samples were snap frozen in liquid nitrogen before transport and were stored at −80 °C prior to analysis.

During this study, 3 different sample cohorts were used: (i) the discovery cohort, (ii) the breed cohort, (iii) the validation cohort. The first discovery cohort consisted of 50 Brown Nick eggs collected at day-7 and 60 eggs at day-8,9,10 and 11 after the start of the incubation. The breed cohort consisted of a total of 146 Brown Nick and 151 Hy-Line eggs, both collected at day-9, and these were used to assess the prediction power of the candidate biomarker in different chicken breeds. Finally, the validation cohort included 143 Brown Nick eggs collected at day-9 and these were analyzed with a targeted LC-APCI-MRM method in order to confirm the identity of the biomarker and its prediction performance.

### 2.3. Genetic gender determination by PCR

To determine the gender of the embryo at the early stages of its development, a PCR analysis was made using the primer 1237L and 1237H (Kahn, St John, & Quinn, 1998). To do so, the embryo toes of each corresponding allantoic fluid sample have been collected and snap frozen in liquid nitrogen. The samples were then subjected to PCR analysis.

### 2.4. Sample preparation

Allantoic samples from the discovery and breed cohort have been deproteinized with methanol. Briefly, 10 μL of the sample has been mixed with 90 μL of cold methanol and then vortexed. After centrifugation at 10,000 rpm at 10°C for 10 min, the supernatant was collected. An equal fraction of each supernatant has been collected for each sample and then mixed together to prepare QC samples which were analyzed at a regular interval between samples.

Prior to targeted analysis, samples from the validation study were spiked with a deuterated internal standard. Briefly, 100 μL of allantoic fluid has been mixed with 100 μL of a solution made of the deuterated internal standard at a concentration of 600 ng/mL in water, using a Tecan Freedom Evo liquid handling platform (Männedorf, Switzerland). QC samples have been prepared by mixing an equal volume of each raw samples and processed in parallel to individual allantoic fluid samples.

### 2.5. RPLC-MS/MSMS

Untargeted metabolite analyses were performed on an ACQUITY UPLC from Waters (Milford, USA) directly coupled to the ESI jet stream nebulizer of an Agilent 6530 series ESI-Q-TOF high-resolution mass spectrometer from Agilent Technologies (Waldbronn, Germany). The separation was performed with a Waters AccQ-Tag Ultra column (1.7 μm, 2.1 mm × 100 mm), thermostated at 60 °C. The LC gradient has been described previously in the analytical chromatography section of our previous work (Laan, Elfrink, Azadi-Chegeni, Dubbelman, Harms, Jacobs,et al., 2021) but with 2% formic acid added to both mobile phases. MS acquisitions were done in ESI positive mode using full scan mode with a mass range from 100-1000 m/z and used a lock mass for improved mass accuracy. MS/MS data were recorded at 5 and 25 eV using an inclusion list of targeted features.

### 2.6. HILIC-MS

HILIC-MS analyses were performed using an Agilent 1260 Infinity LC system directly coupled to a DuoSpray™ ionization source of a Sciex 6600 TripleTOF (Darmstad, Germany). The separation was performed with a Zic-cHILIC column (3 μm, 2.1 mm × 100 mm, 100 Å), purchased from Merck Millipore and thermostated to 25 °C. The gradient is similar to published HILIC chromatography (Laan, Elfrink, Azadi-Chegeni, Dubbelman, Harms, Jacobs,et al., 2021) but used 5mM of ammonium acetate instead of ammonium formate and a flow rate of 0.2 ml/min. MS acquisitions were performed in ESI positive mode using full-scan acquisition mode with a mass range from 50 to 1500 m/z.

### 2.7. LC-APCI-MRM

Targeted analyses were performed on an ACQUITY UPLC from Waters (Milford, USA) directly coupled to the Multimode source of an Agilent 6460 series QqQ mass spectrometer from Agilent Technologies (Waldbronn, Germany). The separation was performed with a Waters AccQ-Tag Ultra column (1.7 μm, 2.1 mm × 100 mm), thermostated at 60 °C. The mobile phases were prepared by adding (A) water and and (B) methanol both with 0.1 % formic acid as modifier. The injection volume was 5 μL, and the flow rate of the mobile phase was set to 0.3 mL/min. Mobile phase B was increased from 0.5 to 30% over 3 min, then was further increased to 99% in 0.1 min. Finally, mobile phase B was brought back to the initial conditions in 0.9 min over a period of 3 min. MS acquisitions were done in APCI positive mode using the MRM acquisition mode. The transitions 164.1 → 119.0 and 168.1→ 119.0 with a collision energy of 12 V and a dwell time of 50 ms were used to monitor 3-[(2-aminoethyl)sulfanyl]butanoic acid and the deuterated version respectively.

### 2.8. Data processing

The raw data from both discovery and breed cohorts were pre-processed using Agilent MassHunter Workstation Software Profinder (Agilent, Version B.06.00, Build 6.0.606.0. The Batch Recursive Feature Extraction was performed on the raw files loaded into the software. The following settings were used, settings not mentioned were as used as default by the software. Molecular feature extraction was performed for peaks between 0 to 8 min, for m/z between 100 and 900 m/z and with a minimum count of 300. Peak binning and RT alignment have been done using a RT window of 1% ± 0.15 min and a mass window = 5.00 ppm ± 2 mDa. The maximum number of features was set to 2000. Recursive feature extraction was made using Find by Ion mode, using matching tolerance of ± 1.5 min for retention time and 35 ppm of symmetric error on the m/z. The peak area extraction of the features was done using Agile algorithm. Duplicate features, artefact features from the data preprocessing, and features present in less than 90% of the samples were manually removed, as well as samples with less than 80% of the total amount of features present.

The raw data from the validation cohort has been processed using Skyline (MacLean, Tomazela, Shulman, Chambers, Finney, Frewen, et al., 2010). Finally, the statistical treatment of the extracted features has been done using MetaboAnalyst 5.0 (Xia, Mandal, Sinelnikov, Broadhurst, & Wishart, 2012). and R packages.

## 3. Results

### 3.1. Allantoic fluid is an accessible and non-damaging fluid for biomarker discovery and routine analysis

The allantois is a hollow-sac structure developed by the embryos and has two main functions. As it is highly vascularized, the allantois *de facto* acts as the lungs of the embryonic chick supplying oxygen for the red blood cells that passed through the eggshell. Its second function is to collect the nitrogenous waste produced by the fetus, thus making the composition of the allantoic fluid close to mammalian urine. It is also interesting to note that having the lowest fluid density, the allantoic fluid is easily accessible after puncturing the shell through the air sac located at the apex of the egg. Consequently, the collection of a reasonable volume, dependent on the development of the fetus, of this biofluid is not detrimental to the fetus and makes it a good source for potential biomarkers (Weissmann, Reitemeier, Hahn, Gottschalk, & Einspanier, 2013).

### 3.2. Untargeted metabolomics approach for the discovery of gender specific biomarkers in allantoic fluid

Metabolomics based biomarker discovery methodologies rely on the detection of a very large variety of compounds in order to maximize the chance of finding interesting hits (Drouin, Pezzatti, Gagnebin, Gonzalez-Ruiz, Schappler, & Rudaz, 2018; Pezzatti, Gonzalez-Ruiz, Codesido, Gagnebin, Joshi, Guillarme, et al., 2019). For this reason, an untargeted liquid chromatography with mass spectrometry (LC-MS) approach was chosen using positive electrospray mode on a high-resolution Q-TOF (Noga, Dane, Shi, Attali, van Aken, Suidgeest, et al., 2012). The discovery cohort comprised 50 eggs for day 7 and 60 at days 8 to 11. In total, 1954 peaks with unique retention time and mass, so called features, were detected in the LC-MS chromatograms with this method. After manual curation of the data, 1390 feature were considered and the gender was modeled as a binary outcome with sampling day and measured features as model predicator. The model was further optimized using backward elimination techniques (Koller & Sahami, 1996)., validated using cross-validation (Steyerberg, Harrell, Borsboom, Eijkemans, Vergouwe, & Habbema, 2001). and evaluated by the percentage of correctly predicted genders, male and female combined together. This approach identified a particular feature (F1558) as the most promising biomarker candidate with a prediction rate up to 91% at day-9 (Figure 1A).

**Figure 1:**
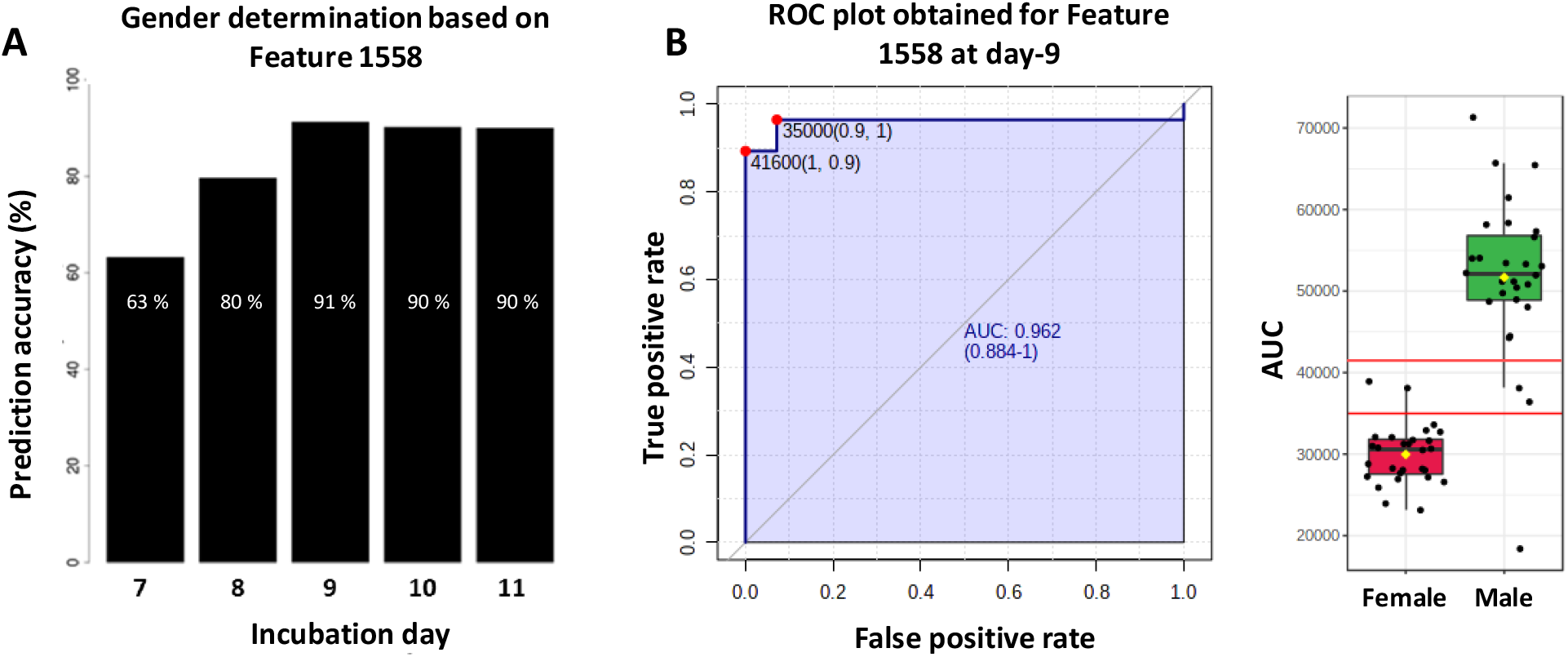
(A) Logistic regression classification model On Single Feature. We evaluated single predictor models for all measured features and observed the best accuracy using Feature 1558 (F1558), which provides >90% accuracy at day-9. (B) ROC plot of F1558 at day-9 and its associated boxplot.

This particular feature presented similar prediction power for days 9 to 11. However, during chicken embryonic development, the neural tube needs approximately 50% of the 21 days of the total gestation duration to turn into a functional brain (Bjornstad, Austdal, Roald, Glover, & Paulsen, 2015; Close, et al., 1997; Mellor & Diesch, 2007). This indicates that chicken embryos can be euthanized up to 11 days after laying, before they can perceive pain. In addition, we found the collection of allantoic fluid from an individual egg during day-9 of the development did not lead to a loss of hatchability of the embryos. For ethical reasons, we choose to focus on day-9 and disregarded later days. A univariate receiver operating characteristic (ROC) analysis for F1558 (shown in Figure 1B) shows a sensitivity of 96.4% (IC_95%_ = 89.3-100%) and a specificity of 92.9% (IC_95%_ = 85.7-100%) using a peak area threshold of 35,000 counts.

The first experiment identified a potential biomarker of gender from day-8 to 11 in H&N Brown Nick chickens. To evaluate the prediction capacity of this feature in different chicken lines, a new experiment was performed with the analysis of a breed-cohort of allantoic fluid coming from Brown Nick chickens and white Hy-Line chickens (146 and 151 samples respectively) collected at day-9. As shown in Figure 2, we did not find any statistical difference of the average biomarker level in the overall population of each chicken line (p_0.05_ = 0.52). When considering brown and white eggs together, we find the endogenous level of the biomarker was significantly different between male and female eggs (p_0.05_ = 1.25 × 10^−44^). It has thus been possible to set a common intensity threshold, independent of the chicken lines, giving a prediction accuracy above 89.5%.

**Figure 2:**
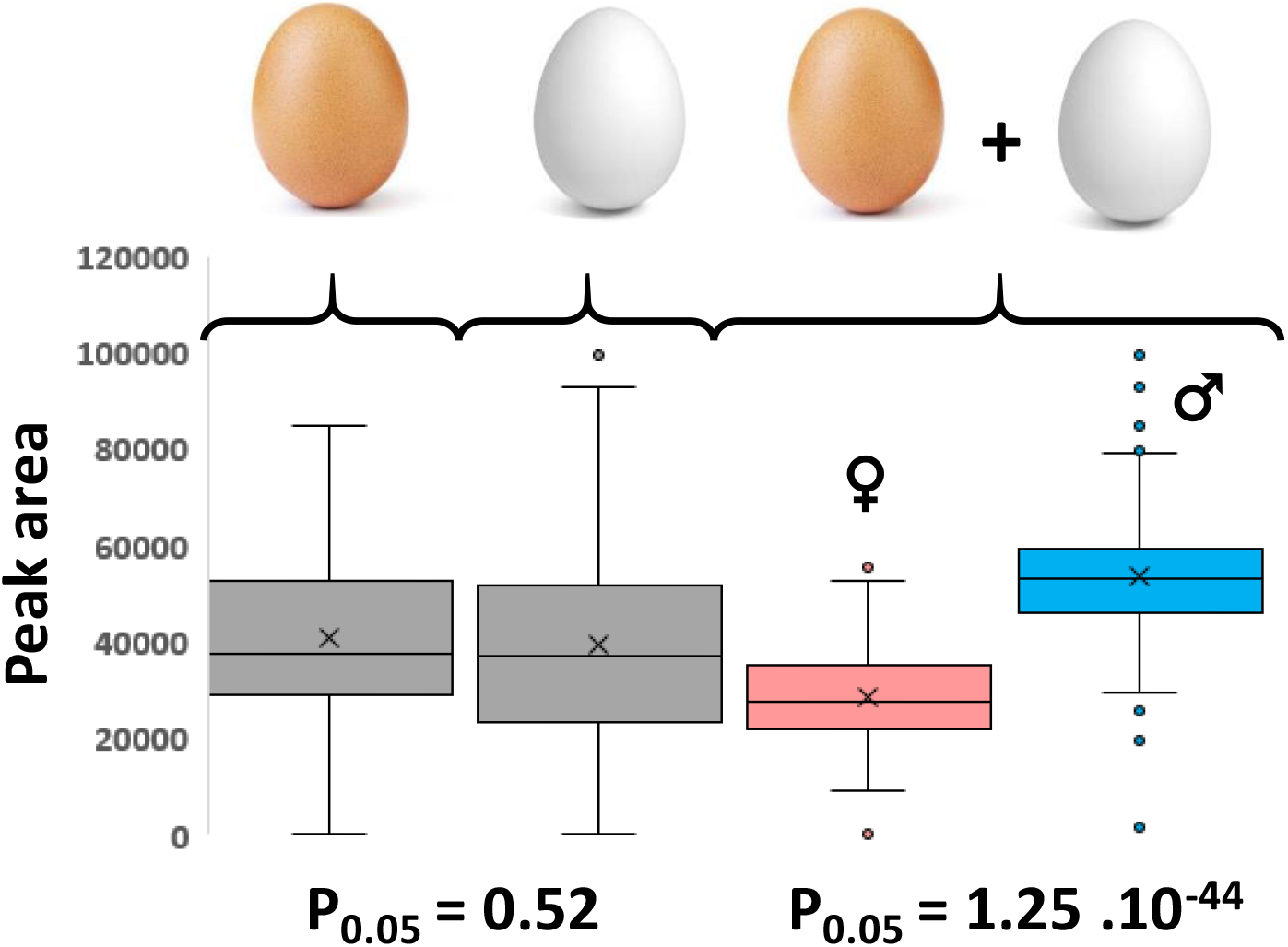
Box-plot comparison of the white and brown eggs populations

These results clearly highlight the strength of the discovered feature for early gender determination of chicken embryos 9 days after the egg-laying regardless of chicken species.

### 3.3. 3-[(2-aminoethyl)sulfanyl]butanoic acid was identified as a gender-specific biomarker

Unambiguous identification of the biomarker requires the comparison of the feature of interest to reference compounds using different characterization approaches to obtain orthogonal information.

Therefore, for confident annotation of an unknown feature, at least 2 independent parameters need to match the reference compound for the feature to be identified, such as liquid chromatography retention time, accurate mass or mass spectrometry fragmentation pattern (MSMS spectra) (Rochat, 2017; Sumner, Amberg, Barrett, Beale, Beger, Daykin, et al., 2007).

The exact mass of the protonated form of F1558 has been accurately measured as m/z 164.0739. This mass was assigned by Agilent MassHunter as corresponding to the chemical formula C_6_H_13_NO_2_S which was confirmed by the isotopic distribution showing the specific pattern of sulfur atoms (Figure 3A). In order to identify the biomarker, we undertook the identification of its MSMS fragments obtained after RPLC-MSMS of a pooled allantoic sample (Figure 3B). Based on the fragmentation pattern, we proposed different structural isomers, whereby S-propyl-L-cysteine, S-(2-carboxypropyl)-cysteamine, and 3-[(2-aminoethyl)sulfanyl] butanoic acid were the most likely to produce the same fragments (Figure 3C). Since they were not commercially available, these compounds were synthesized on our request.

**Figure 3:**
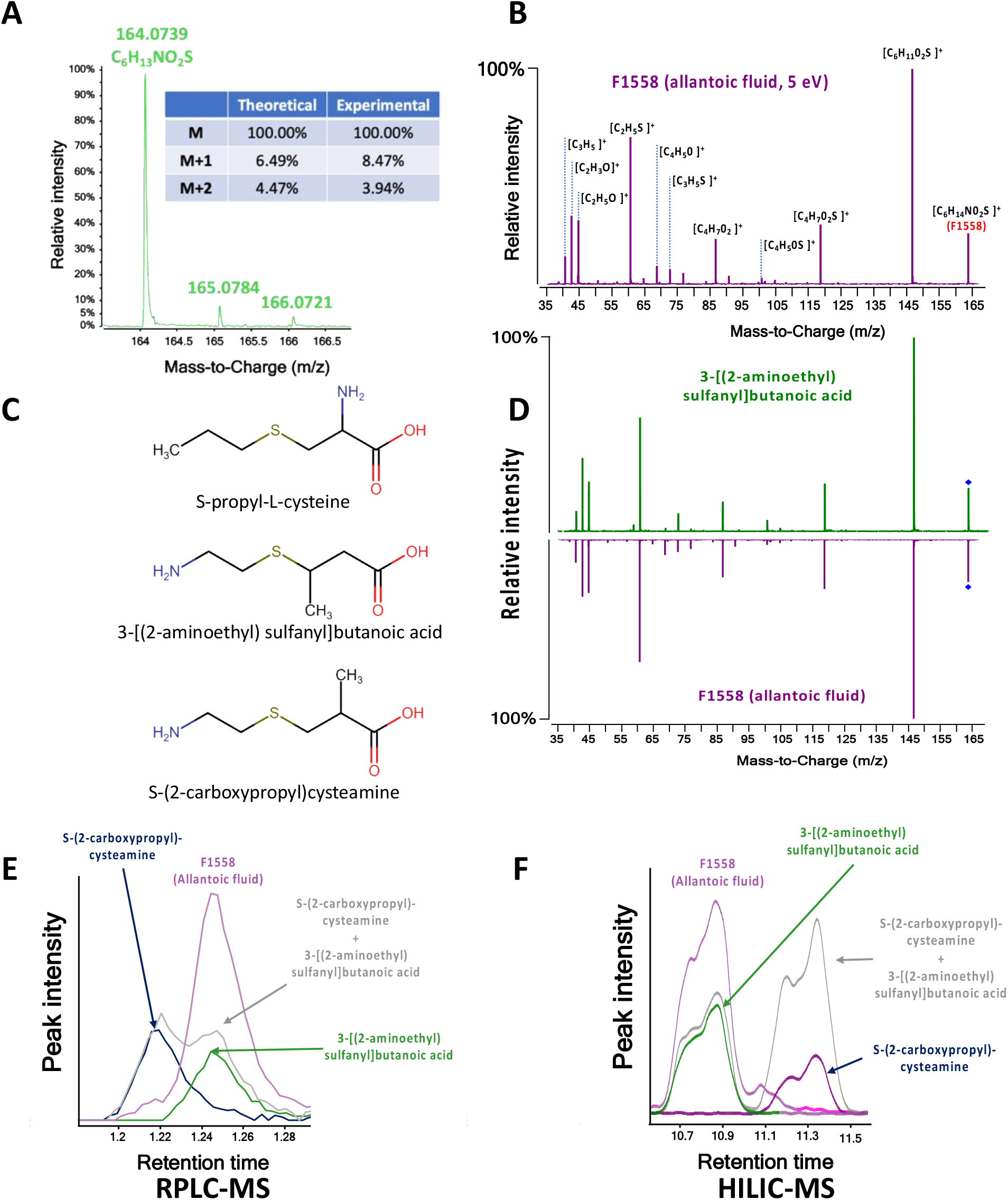
Identification process of F1558 (A: determination of the elemental composition, B: MSMS spectra of F1558 and elemental composition of fragments, C: possible structures, D: comparison of fragmentation pattern of F1558 and 3-[(2-aminoethyl)sulfanyl] butanoic acid, E-F: retention profile of F1558, S-propyl-L-cysteine, S-(2-carboxypropyl)-cysteamine and 3-[(2-aminoethyl)sulfanyl] butanoic acid in RPLC and HILIC mode respectively)

The fragmentation pattern of the biomarker F1558 was then compared to the MSMS spectra of the three candidate compounds. It quickly appeared that 3-[(2-aminoethyl)sulfanyl] butanoic acid possessed 11 common fragments with F1558 (Figure 3D) whereas the two other compounds only had 2 (Table S1). Additional confirmation of the identity was obtained by comparing retention time profiles in two orthogonal chromatography methods. After injection in RPLC mode, S-propyl-cysteine was quickly discarded from the list since its retention time (RT) differed by more than one minute from F1558 (data not shown). The RT obtained for 3-[(2-aminoethyl)sulfanyl] butanoic acid and S-(2-carboxypropyl)-cysteamine were slightly different, and two peaks were clearly observed when co-injected (Figure 3E). We found the RT of 3-[(2-aminoethyl)sulfanyl] butanoic acid was exactly the same as F1558. Second, we undertook a HILIC comparison. As shown in Figure 3F, in this mode, 3-[(2-aminoethyl)sulfanyl] butanoic acid and S-(2-carboxypropyl)-cysteamine gave two baseline separated peaks. The peak shape and the RT of 3-[(2-aminoethyl)sulfanyl] butanoic acid were strictly identical to F1558. Therefore, combining all these orthogonal data together, we were able to unequivocally identify 3-[(2-aminoethyl)sulfanyl]butanoic acid (ASBA) as being the feature of interest F1558.

### 3.4. ASBA shows high sensitivity and specificity for gender prediction

Based on a small discovery cohort using untargeted metabolomics profiling of allantoic fluid at day-9, we identified ASBA as a potential biomarker for *in ovo* gender prediction of chicken eggs. In order to validate our hypothesis and to determine the prediction power of ASBA, we investigated a validation cohort including 143 allantoic fluid samples collected from Brown Nick eggs at day-9. Untargeted metabolomics methods are known to have reduced accuracy due to matrix effect and low peak abundance. For this reason, we decided to validate the candidate biomarker using a targeted LC-APCI-MRM method and using the deuterated version of the metabolite as internal standard.

As highlighted in Figure 4 by the increase of the AUC of the ROC plot, the improvement of the analytical method led to an increase of the prediction accuracy, leading to a total separation of the male embryos with a sensitivity of 100% (IC_95%_ = 100-100%) and a specificity of 96.2% (IC_95%_ = 92.3-100%) for a peak area ratio threshold established at 0.342.

**Figure 4:**
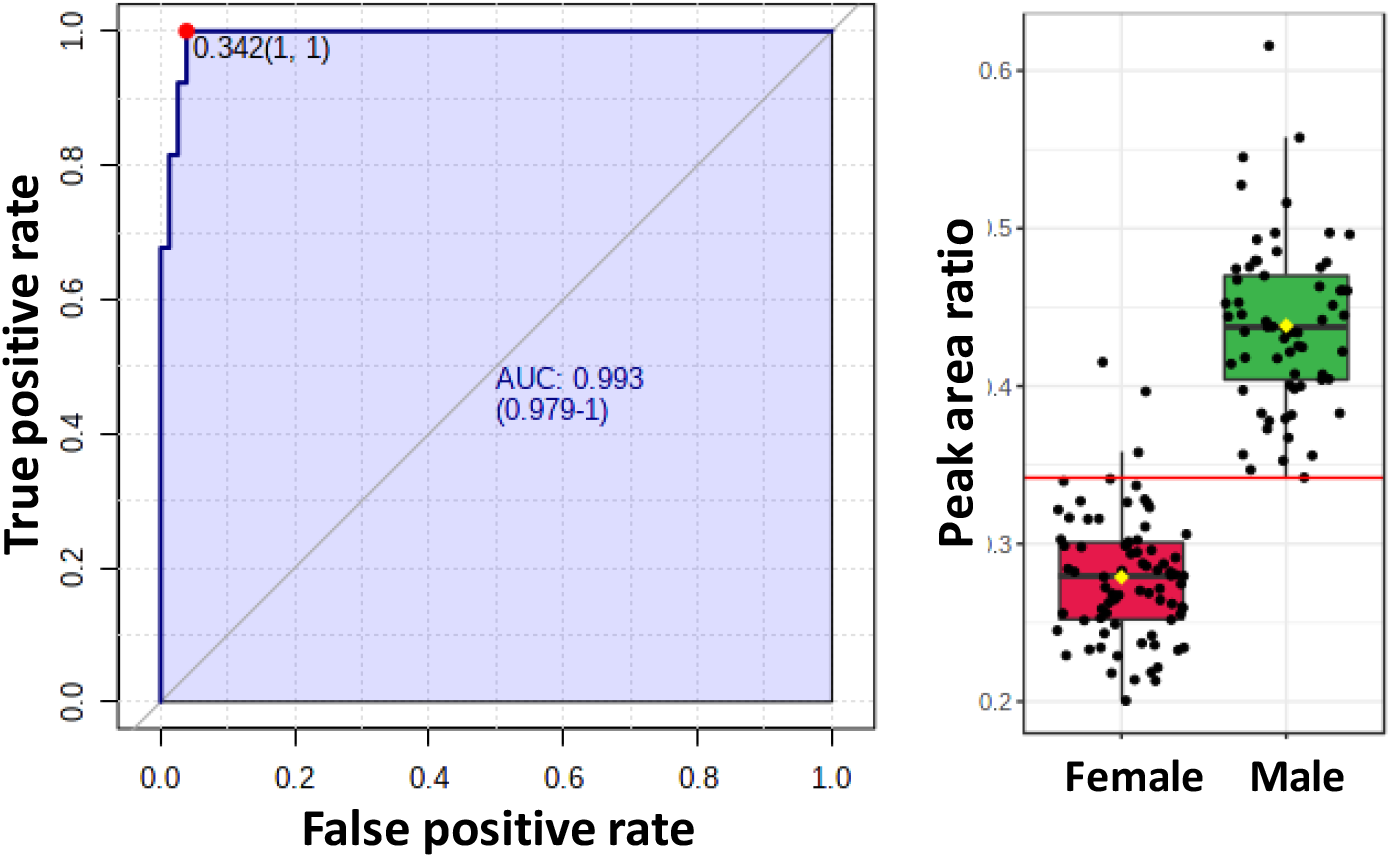
ROC and box plots obtained with targeted LC-APCI-MRM analysis.

To study the prediction performance of the model, we so far have used a male/female cut-off determined *a posteriori* and based on true gender. Obviously, this is not possible in routine analysis and we attempt to set a fixed cut-off based on QC samples. Indeed, the theoretical distribution of males and females in a population is a Gaussian curve with an average close to 50%. Therefore, we used the peak area ratio of QC samples (i.e. 0.345) prepared from equal volumes of every sample as threshold for the gender determination. Using this approach, the model was able to predict the male gender with a sensitivity of 98.5% (IC_95%_ = 96.1-100%) and a specificity of 96.2% IC_95%_ = 92.3-99.4%).

## 4. Discussion

With the changing mindset and the rapid evolution of legislation, finding alternatives to the culling of day-old male chicks has become a societal issue. In this context, finding a method to *in ovo* discriminate male and female chickens, before the full development of the central nervous system appears to be a humane solution to this problem. Applying untargeted LC-MS approaches on limited sample cohorts, we were able to identify a candidate biomarker feature, and with it able to accurately predict the gender of chicken embryos from day-8 to 11 after egg-laying. We have also demonstrated the prediction accuracy of this candidate biomarker across two different chicken breeds. After its identification, we have been able to confirm that 3-[(2-aminoethyl)sulfanyl]butanoic acid (ASBA) is a powerful biomarker for early gender screening. Surprisingly, this is the first report of this gender-specific biomarker in chickens or any living species. Finally, we have validated this biomarker using a targeted LC-APCI-MRM method on more than 140 samples and we showed by only diluting the raw allantoic fluid with the corresponding deuterated compound, we were able to improve the prediction power of the method to a sensitivity of 100% and a specificity of 96.2%. Nevertheless, a posteriori threshold is not possible in routine analysis. To circumvent this issue, we have shown the approach based on a pooled QC sample and the theoretical distribution of the gender in a population was able to reach a gender prediction accuracy of 97.2 %. Beyond the discovery of ASBA as a powerful early biomarker of chick gender, we successfully developed a high throughput gender screening workflow able to gender screen more than 2000 eggs per our with a prediction accuracy above 95%. This method, using automation and the acoustic droplet ejection hyphenated with mass spectrometry (ADE-MS) is the object of a separate research article (ref) and is now commercialized by In Ovo (Leiden, the Netherlands).

## Acknowledgment

In Ovo Holding B.V. acknowledges support from the Dutch Ministry of Economic Affairs. Alida Kindt-Dunjko is gratefully acknowledged for her support with the data processing.

## CrediT authorship contribution statement

Nicolas DROUIN: Conceptualization, Methodology, Formal analysis, Validation, Investigation, Data curation, Writing – original draft, Writing – Review & Editing, Visualization

Hyung Lim ELFRINK: Conceptualization, Methodology, Formal analysis, Validation, Investigation, Data curation, Writing – original draft, Writing – Review & Editing, Visualization

Slavik KOVAL: Data curation

Amy HARMS: Conceptualization, Methodology, Resources, Data curation, Supervision, Writing – original draft, Writing – Review & Editing, Visualization, Project administration, Funding acquisition

Will STUTTERHEIM: Resources, Project administration, Funding acquisition Wouter BRUINS: Resources, Project administration, Funding acquisition

Thomas HANKEMEIER: Conceptualization, Methodology, Resources, Data curation, Supervision, Writing – original draft, Writing – Review & Editing, Visualization, Project administration, Funding acquisition

